# An improved CRISPR/dCas9 interference tool for neuronal gene suppression

**DOI:** 10.1101/2020.05.26.116822

**Authors:** Corey G. Duke, Svitlana V. Bach, Jasmin S. Revanna, Faraz A. Sultan, Nicholas T. Southern, M. Natalie Davis, Nancy V.N. Carullo, Allison J. Bauman, Robert A. Phillips, Jeremy J. Day

**Affiliations:** Department of Neurobiology & Evelyn F. McKnight Brain Institute, University of Alabama at Birmingham, Birmingham, AL 35294, USA

**Author notes:** Corresponding author *Correspondence to Jeremy Day* ( | day-lab.org | @DayLabUAB).

## Abstract

The expression of genetic material governs brain development, differentiation, and function, and targeted manipulation of gene expression is required to understand contributions of gene function to health and disease states. Although recent improvements in CRISPR/dCas9 interference (CRISPRi) technology have enabled targeted transcriptional repression at selected genomic sites, integrating these techniques for use in non-dividing neuronal systems remains challenging. Previously, we optimized a dual lentivirus expression system to express CRISPR-based activation machinery in post-mitotic neurons. Here we used a similar strategy to adapt an improved dCas9-KRAB-MeCP2 repression system for robust transcriptional inhibition in neurons. We find that lentiviral delivery of a dCas9-KRAB-MeCP2 construct driven by the neuron-selective promoter human synapsin 1 enabled transgene expression in primary rat neurons. Next, we demonstrate transcriptional repression using CRISPR sgRNAs targeting diverse gene promoters, and show superiority of this system in neurons compared to existing RNA interference methods for robust transcript specific manipulation at the complex Brain-derived neurotrophic factor (*Bdnf*) gene. Our findings advance this improved CRISPRi technology for use in neuronal systems for the first time, potentially enabling improved ability to manipulate gene expression states in the nervous system.

---

BRAIN FUNCTION and development relies on tightly coordinated transcriptional programs, and dysregulated gene expression patterns are linked to many neurological and psychological disorders^1–14^. Understanding the functional relevance and contributions of divergent expression states requires the ability to manipulate gene expression in a targeted and specific manor. A classic approach is to delete an individual candidate gene implicated in a biological process or disease state and characterize the resulting phenotype. Although powerful, this approach often carries significant drawbacks^15^, such as those arising from genetic background^16^, genetic compensation unrelated to the loss of the targeted gene’s protein function^17^, an inability to assess genes for which a knockout is lethal, and technical challenges in achieving homogenous knockouts at all loci due to ploidy^18^. It is also often difficult to distinguish the functional role of a gene of interest in behavior from its role in development^15^ due to the irreversibility of genetic manipulations^18^. Further, this approach suffers from an inherent lack of specificity in cases where the entire genomic locus is perturbed rather a specific transcript isoform. RNA interference (RNAi) based methods of gene suppression can overcome many of these challenges, but also possess extensive sequence-dependent and sequence-independent off-target effects^19–25^. These limitations can make interpretations of knockout and RNAi studies challenging, and highlight the need for more robust strategies to manipulate gene expression^21–27^.

Advances in CRISPR/Cas9 technology have revolutionized functional investigations of gene expression^28–31^. Cas9, an RNA-directed endonuclease, can be localized to selected genomic sites with a single guide RNA (sgRNA) complementary to the DNA location of interest, provided a protospacer adjacent motif (PAM) sequence is nearby (5’-NGG-3’ for the widely used *Streptococcus pyogenes* Cas9). Once trafficked, it will induce a double stranded break andcantherebybeusedtocreatetargetedpermanentgenomic alterations^32^. Mutation of the catalytic domain of Cas9 to generate nuclease-dead Cas9 (dCas9) preserves its homing function and also enables a diverse array of applications^31,33–35^. For example, dCas9 targeted to the transcriptional start sites of genes physically impedes the transcription process without altering the DNA sequence itself, a process termed CRISPR interference (CRISPRi)^36^. This strategy can be preferable to Cas9 approaches when assaying gene expression due to the avoidance of cellular toxicity arising from the induced double-stranded DNA breaks, the increased specificity of altering transcription while preserving the genetic structure, and the potential reversibility of the resultant transcriptional modifications^36–42^. Subsequent efforts have improved the transcriptional silencing function of CRISPRi via fusion of potent transcriptional inhibitors to dCas9^33,38^, such as a Krüppel-associated box (KRAB) repressor domain. While this approach has been adopted for targeted as well as multiplexed gene silencing in the nervous system^43^, these approaches have not led to consistent gene silencing at other targets^44^, highlighting the need for continued optimization.

A recent unbiased screen comparing multiple dCas9 fusion systems demonstrated robust and selective gene repression via fusion of a dimer containing a KRAB repressor domain in addition to methyl-CpG binding protein 2 (MeCP2). CRISPRi with dCas9-KRAB-MeCP2 resulted in significant improvement over existing dCas9 interference approaches, as well as gene knockdown using classic RNAi technology^45^. However, the adoption of this technology for use in neuronal systems requires surpassing significant barriers due to difficulties in transgene delivery and nuclear localization in post-mitotic neurons^44,46,47^. Recently, we established an approach for CRISPR activation (CRISPRa) to achieve specific, robust, and multiplexable gene induction in neurons both *in vitro* and *in vivo* by engineering an optimized dual-lentivirus technique^44^. Here, we apply these advances to translate this second-generation dCas9-KRAB-MeCP2 CRISPRi technology for functional use in neuronal systems.

## Results

### ASYN-drivendCas9-KRAB-MeCP2 construct suppresses gene expression at a luciferase reporter in HEK293T cells

As highlighted in previous studies, catalytically inactivated Cas9 (dCas9) fusion systems recruiting KRAB and MeCP2 domains provide improved gene-specific transcriptional knockdown (CRISPRi) in mammalian cell lines^45,54^ (**Figure 1a**). To leverage this second-generation construct for efficient use in neuronal systems, we adopted a dual lentiviral approach segregating the dCas9-KRAB-MeCP2 and the sgRNA onto separate vectors (**Figure 1b**). This approach enables efficient packaging of the large dCas9-KRAB-MeCP2 fusion with the robust neuron-selective human synapsin 1 promoter (SYN) and provides the flexibility to readily exchange and combine sgRNA scaffold constructs^44^. To confirm functionality of the dCas9-KRAB-MeCP2 construct, we engineered a luciferase reporter system containing firefly luciferase positioned downstream of the truncated rat *Fos* promoter in HEK293T cells. Transfecting sgRNAs designed to target the adapted rat *Fos* promoter region produced robust suppression of luciferase signal relative to control sgRNAs targeted to the bacterial gene *lacZ* (a sequence that does not exist in the mammalian genome) (**Figure 1c**), demonstrating that the adapted SYN-dCas9-KRAB-MeCP2 fusion system produces efficient targeted gene suppression that translates into a loss of functional protein.

**Figure 1.**
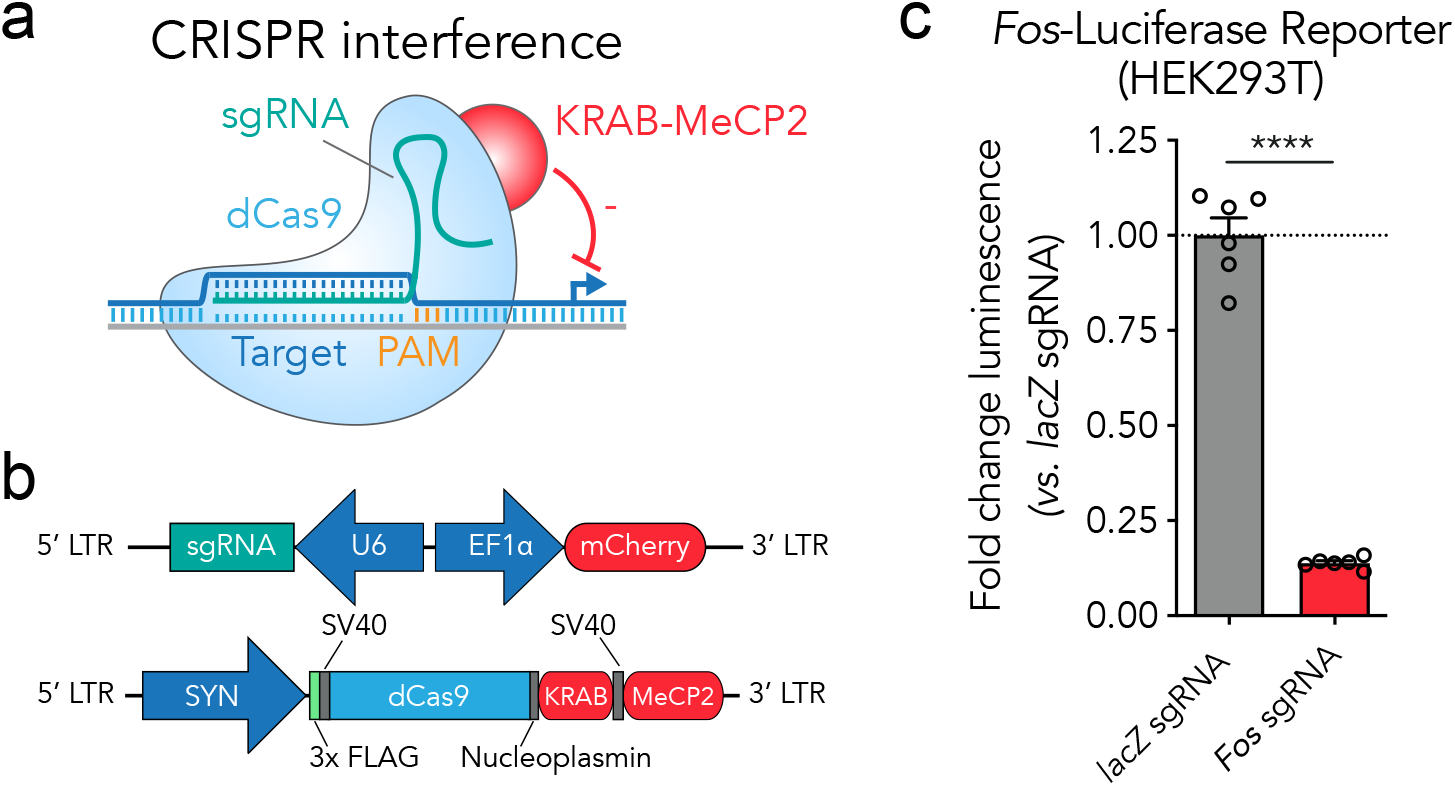
A SYN-driven dCas9-KRAB-MeCP2 construct suppresses gene expression at a luciferase reporter in HEK293T cells. (**a**) Illustration of the dCas9-KRAB-MeCP2 suppression strategy. An sgRNA with a spacer complementary to the targeted genomic site adjacent to a PAM motif directs the dCas9-KRAB-MeCP2 transcriptional suppresser to targeted genetic loci. (**b**) The dual vector lentiviral construct designs. The U6 polymerase 3 promoter drives expression of the sgRNA which can be adapted to target specific genetic loci. The EF1α promoter drives expression of mCherry, useful for rapid assessment of lentiviral expression. The second construct contains the dCas9-KRAB-MeCP2 fusion driven by the neuron-selective SYN promoter. This fusion contains a FLAG epitope so that construct expression can be visualized readily by immunocytochemistry. (**c**) Luciferase assay confirms targeted gene suppression by the dCas9-KRAB-MeCP2 system in HEK293T cells relative to non-targeted (*lacZ*) sgRNA controls (*n* = 6, unpaired *t*-test, *t*_(10)_ = 18.68, *p* < 0.0001). All data are expressed as mean ± s.e.m. Individual comparisons; \*\*\*\**p* < 0.0001.

### dCas9-KRAB-MeCP2 is capable of strong gene suppression at multiple genes in primary neuronal cultures

To examine functionality of the adapted dCas9-KRAB-MeCP2 system in neuronal systems, plasmid constructs were packaged into high-titer lentiviruses (5.47*10^10^ to 1.91*10^11^ genomic copies per mL) and transduced into primary rat neuronal cultures (**Figure 2**). The dCas9-KRAB-MeCP2 fusion was engineered to contain the FLAG epitope, which readily enables immunocytochemistry (ICC) visualization studies. ICC against the FLAG epitope revealed strong transduction efficiency, neuronal expression, and nuclear localization of the dCas9-KRAB-MeCP2 fusion in rat primary striatal cultures (**Figure 2a**). To confirm sgRNA-targeted repression and better understand the lentiviral load required when utilizing this system in neurons, we varied the lentiviral genomic copies delivered per cell while targeting the immediate early gene *Fosb* in primary rat hippocampal cultures (**Figure 2b**). A two-way ANOVA revealed a significant main effect of sgRNA for the *Fosb* groups relative to *lacZ* controls, suggesting effective gene silencing. In contrast, there was no main effect of lentiviral genome copy number, indicating that 1000 genome copies per cell was sufficient for primary neuronal culture experiments. To examine the capability of this system to downregulate genes of diverse functional classes, we developed and employed sgRNAs to traffic the system to the lysine methyltransferase *Kmt2b*, the extracellular matrix protein *Reln*, and the signaling neuropeptide *Npy* (**Figure 2c**). Comparison of gene expression with gene-specific RT-qPCR primer sets revealed significant decreases in gene expression at all targets compared to *lacZ* control sgRNAs. These results confirm that robust targeted gene repression can be achieved using the dCas9-KRAB-MeCP2 system across diverse gene targets in neuronal systems.

**Figure 2.**
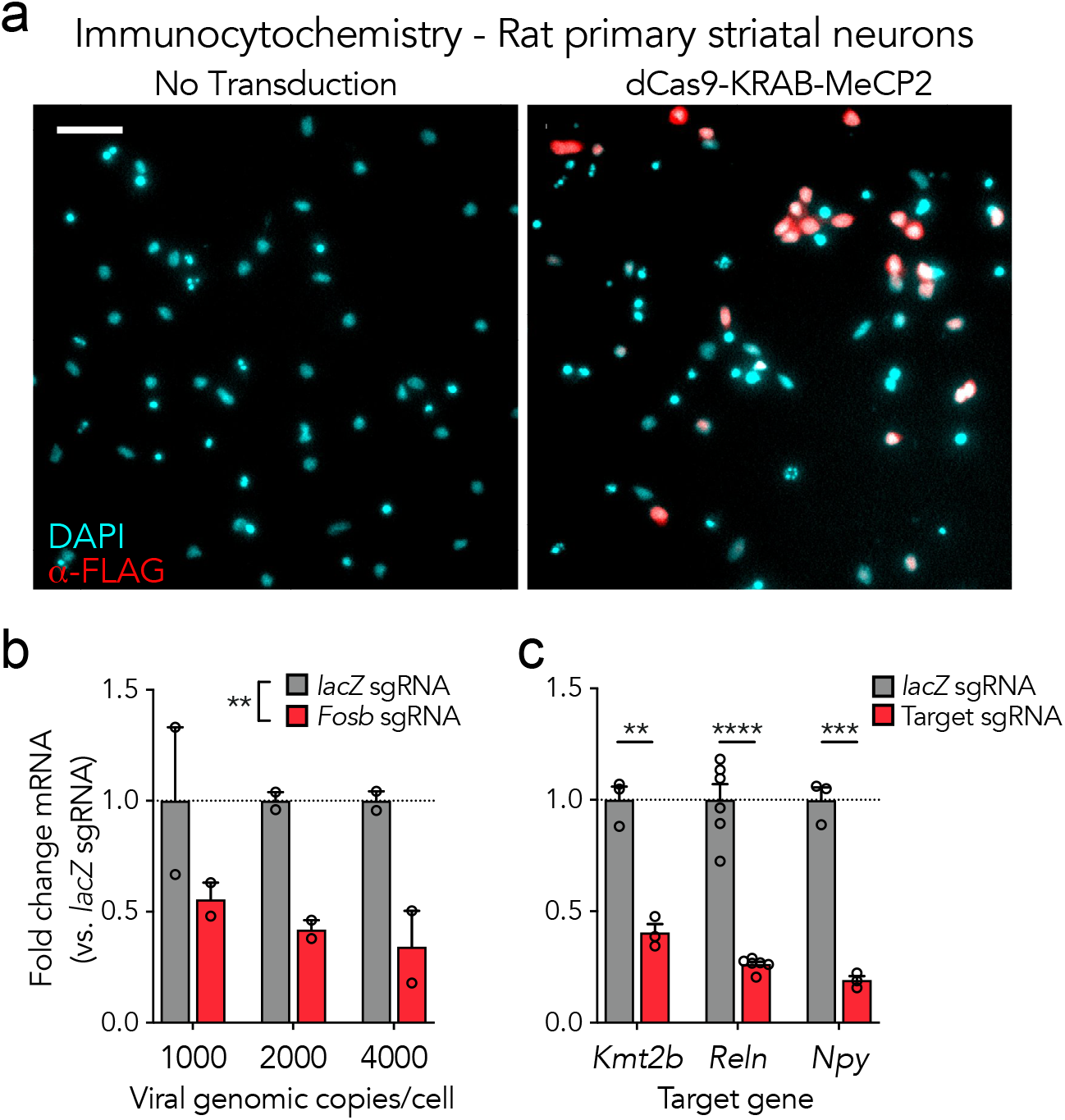
dCas9-KRAB-MeCP2 is capable of strong gene suppression at multiplegenesinprimaryneuronalcultures. (**a**) Immunocytochemistry demonstrating expression of the lentiviral SYN-driven dCas9-KRAB-MeCP2 construct in primary rat striatal cultures. Scale bar = 200μm. (**b**) dCas9-KRAB-MeCP2 induces targeted gene suppression at *Fosb* relative to *lacZ* sgRNA controls in primary rat hippocampal cultures as revealed by RT-qPCR. There was no main effect of viral genome copies per cell (*n* = 2, two-way ANOVA; sgRNA *F*_(1, 6)_ = 19.17, *p* = 0.0047; viral genomic copies per cell *F*_(2, 6)_ = 0.2380, *p* = 0.7953, interaction *F*_(2, 6)_ = 0.2380, *p* = 0.7953). (**c**) RT-qPCR demonstrates targeted gene suppression in striatal cultures relative to non-targeted controls across multiple genes (*n* = 3-6, unpaired *t* test; *Kmt2b t*_(4)_ = 8.373, *p* = 0.0011; *Reln t*_(10)_ = 10.34, *p* < .0001; *Npy t*_(4)_ = 13.79, *p* = 0.0002). All data are expressed as mean ± s.e.m. Individual comparisons, \*\**p* < 0.01, \*\*\**p* < 0.001, \*\*\*\**p* < 0.0001.

### dCas9-KRAB-MeCP2 induces targeted transcript-specific Bdnf gene repression in primary rat hippocampal cultures

Gene expression in neuronal systems is complex, with many genes utilizing alternative splicing in critical ways to control development and synaptic plasticity^55–58^. One example of this complexity occurs at the brain-derived neurotrophic factor gene *Bdnf*, which uses divergent non-coding exons (*1-Xa*) combined with a single coding exon (*IX*) in multiple alternatively spliced forms produced from distinct gene promoters (**Figure 3a**)^59^. The most highly expressed transcripts employ either the non-coding exon *I* or *IV*, and the specific functionality of this transcript heterogeneity remains an area of active investigation^44,60–62^. We designed short hairpin RNAs (shRNAs) and CRISPRi sgRNAs to compare these targeted transcript suppression methods at the *Bdnf I* and *Bdnf IV* individual transcripts. shRNA targeting of the *Bdnf I* transcript was unsuccessful at decreasing its expression, but instead resulted in an unintended decrease in the expression of *Bdnf IV* (**Figure 3b**). *Bdnf IV* shRNA successfully repressed *Bdnf IV* expression and resulted in a surprising increase of *Bdnf I* transcript expression. Both shRNAs failed to reduce total coding *Bdnf* levels as measured by RT-qPCR primers located in the *Bdnf IX* coding region. Taken together, these findings suggest that while capable of some transcript-specific knockdown, these shRNAs produced only modest effects at the targeted transcripts, a feature commonly observed in RNAi strategies^18,37^. To examine if a dCas9-KRAB-MeCP2 strategy could improve transcript-specific knockdown levels, we next employed individual sgRNAs designed to target regions upstream of the *Bdnf I* and *Bdnf IV* exons in hippocampal neurons (**Figure 3c**). Strikingly, dCas9-KRAB-MeCP2 targeting induced a 96% transcriptional knockdown at *Bdnf I* and a 92% knockdown at *Bdnf IV* relative to *lacZ* controls. Interestingly, targeting the *Bdnf IV* upstream region did not affect levels of *Bdnf I*, but targeting the *Bdnf I* upstream region resulted in a 21.9% reduction of *Bdnf IV* levels. Targeting either region resulted in a significant reduction in *Bdnf IX* levels (35.8% for the *Bdnf I* sgRNA and 42.7% for the *Bdnf IV* sgRNA). Taken together, these findings indicate that the dCas9-KRAB-MeCP2 system is capable of transcript-selective knockdown and can outperform widely utilized RNAi knockdown methods in neuronal systems.

**Figure 3.**
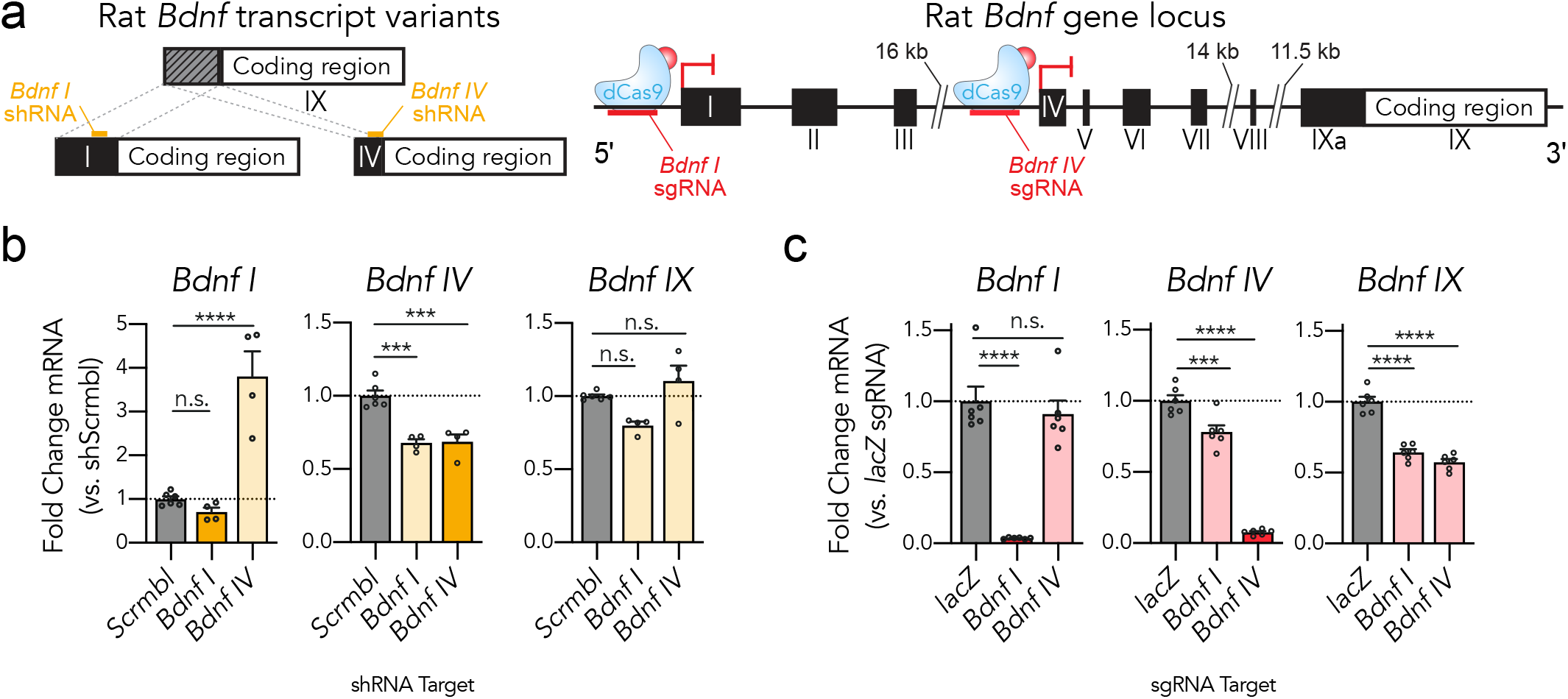
dCas9-KRAB-MeCP2 induces targeted transcript specific *Bdnf* gene repression in primary rat hippocampal cultures. (**a**) Complex structure of the rat *Bdnf* gene produces spliced transcripts from multiple promoter regions using non-coding exons (*I-IXa*) and a common coding exon (*IX*). shRNAs (*left*) or sgRNAs (*right*) can be designed for targeted transcript suppression. (**b**) shRNA targeting the *Bdnf I* variant did not significantly decrease its expression as assessed by RT-qPCR, while shRNA designed against the *Bdnf IV* variant increased expression of the *Bdnf I* variant (*left*, *Bdnf I* RT-qPCR: *n* = 4-6, one-way ANOVA, *F*_(2, 11)_ = 32.55, *p* < 0.0001. *Middle*, *Bdnf IV* RT-qPCR: *n* = 4-6, one-way ANOVA, *F*_(2, 11)_ = 25.19, *p* < 0.0001). Targeting either variant did not alter the expression levels of total *Bdnf* as measured using qPCR primers designed for the *Bdnf IX* common coding region (*right*, *Bdnf IX* RT-qPCR: *n* = 4-6, one-way ANOVA, *F*_(2, 11)_ = 7.266, *p* = 0.0097). (**c**) sgRNAs designed upstream of the *Bdnf I* and *Bdnf IV* exons recruit dCas9-KRAB-MeCP2 to induce transcript specific gene repression of the *Bdnf I* and *Bdnf IV* transcript variants as assessed by RT-qPCR. Both sgRNAs resulted in a significant decrease in total *Bdnf IX* expression (*left*, *Bdnf I* RT-qPCR: *n* = 6, one-way ANOVA, *F*_(2, 15)_ = 41.87, *p* < 0.0001. *Middle*, *Bdnf IV* RT-qPCR: *n* = 6, one-way ANOVA, *F*_(2, 15)_ = 181.9, *p* < 0.0001. *Right*, *Bdnf IX* RT-qPCR: *n* = 6, one-way ANOVA, *F*_(2, 15)_ = 74.56, *p* < 0.0001). Tukey’s multiple comparisons test was used for individual comparisons. All data are expressed as mean ± s.e.m. Individual comparisons; \*\*\**p* < 0.001, \*\*\*\**p* < 0.0001.

## Discussion

Here, we describe an adapted CRISPRi system in which a recently optimized dCas9-KRAB-MeCP2 transcriptional repressor is driven by the human synapsin promoter. Lentiviral expression of sgRNAs targeting this system to unique gene promoters produced robust knockdown at multiple gene targets in two neuronal culture platforms. This optimized CRISPRi system also enabled transcript-specific manipulations at the *Bdnf* gene locus, revealing efficient silencing of a complex target relevant to learning, memory, and neuropsychiatric disease^59,63–70^.

It is worth highlighting the extent of gene depletion achievedbyasinglesgRNAintheseexperiments. Forinstance, near knockout levels of transcript-selective suppression were achieved at *Bdnf* without disrupting the genetic locus itself (96% at *Bdnf I* and 92% at *Bdnf IV*). However, not all targeted gene transcripts investigated were suppressed to the same level. The majority of the sgRNAs utilized here were originally designed for transcriptional activation by placing them in the promoter region upstream of gene transcription start sites (TSS)^44^, not downstream of the TSS as is commonly suggested for CRISPRi^71^. Enhanced silencing strength could likely be achieved by using sgRNAs designed specifically to interfere with transcription elongation and transcription factor binding^71^, or by multiplexing multiple sgRNAs together to target genes of interest^45^. However, our observation that upstream TSS sgRNAs could be useful for CRISPRi may also reflect an advantage of this optimized system, permitting bidirectional gene modulation (i.e., CRISPRa and CRISPRi) using the same sgRNA in parallel experiments.

Recently, we harnessed a neuron-optimized dCas9 transcriptional activator system to target and induce 16 genes simultaneously to mimic the genetic signature of acute dopamine receptor activation^72^. Such strategies enable an unprecedented ability to investigate larger transcriptional networks rather than relying on studies of individual genes. Likewise, recent studies have devoted additional attention to modulation of non-coding RNA and cis-regulatory element function using dCas9 strategies^73,74^. The adapted CRISPRi dCas9-KRAB-MeCP2 system described here is compatible with these approaches and can easily be harnessed for expanded gene silencing via the use of multiplexed sgRNA expression cassettes, targeting non-coding RNA loci, or trafficking the system to regulatory regions of interest.

The system described here uses a neuron-selective promoter, which will enable selective targeting of neurons when applied for *in vivo* uses^44^. Given the cellular diversity of the nervous system, future extensions of this technology to target individual cell populations through combination with cre-recombinase systems is of high interest. Recently, the lentiviral delivery of the first generation dCas9-KRAB construct was employed to selectively modify expression states in either glutamatergic or GABAergic neurons by using mouse CaMKIIα and VGAT selective promoters^43^. Our construct could be adapted similarly for targeted transcriptional manipulations in specific neuronal populations. This report also demonstrated that the first generation dCas9-KRAB system provided more robust silencing as compared to shRNAs at multiple genes in primary neurons^43^. Thus, our results add to accumulating evidence supporting increased efficiency of CRISPRi over RNAi in achieving robust gene repression^18,45^.

As this system does not permanently alter the genetic loci of interest when suppressing transcription, it is possible that transient silencing can be achieved. This has broad appeal for investigations of transient experience-dependent transcription, as well as for critical windows of development. Through combination with drug-^75^ or light-inducible^76–78^ techniques, these approaches may enable temporally specific manipulation that more thoroughly mimics endogenous gene expression patterns. Finally, given the increased transcriptional complexity and alternative splicing that occurs at genes highly expressed in brain tissue^56–58,79,80^, this technology extends our ability to dissect the distinct function of unique transcripts. The need for such technology is ever more apparent as the association of alternative spliced transcripts with neuropsychiatric disease becomes increasingly appreciated^55,58,79,81–85^. In summary, our results demonstrate robust gene silencing in neurons using a second-generation CRIPSRi system, and suggest that continued development of this approach will enable novel experimental strategies in neuronal systems.

## Methods

### Animals

All experiments were performed in accordance with the University of Alabama at Birmingham Institutional Animal Care and Use Committee. Sprague-Dawley timed pregnant rat dams were purchased from Charles River Laboratories. Dams were individually housed until embryonic day 18 for cell culture harvest in an AAALAC-approved animal care facility on a 12-hour light/dark cycle with *ad libitum* food and water.

### Primary Rat Brain Cultures

Primary rat cell cultures were generated from embryonic day 18 (E18) rat striatal and hippocampal tissue, as described previously^44,48–50^. Briefly, cell culture plates (Denville Scientific Inc.) were coated overnight with poly-L-lysine (Sigma-Aldrich; 50 μg/ml) supplemented with laminin (Sigma-Aldrich; 7.5μg/mL) and rinsed with diH_2_O. Dissected cortical or hippocampal tissue was incubated with papain (Worthington LK003178) for 25 min at 37°C. After rinsing in complete Neurobasal media (Neurobasal Medium (Gibco; #21103049), supplemented with B27 (Gibco; #17504044, 1× concentration) and L-glutamine (Gibco; # 25030149, 0.5mM), a single cell suspension was prepared by sequential trituration through large to small fire-polished Pasteur pipettes and filtered through a 100 μm cell strainer (Fisher Scientific). Cells were pelleted, re-suspended in fresh media, counted, and seeded to a density of 125,000 cells per well on 24-well culture plates (65,000 cells/cm^2^). Cells were grown in complete Neurobasal media for 11 days *in vitro* (DIV 11) in a humidified CO_2_ (5%) incubator at 37°C with half media changes at DIV 1 and 10. Lentiviral transduction occurred on either DIV 4 or 5 when 330μl of culture media was removed from each culture well and lentivirus was delivered for an incubation period of 8-12hr. Following this transduction period, 600μl of fresh complete Neurobasal media wash occurred before replacement with a mixture of 300μl of fresh complete Neurobasal and 300μl of the culture media removed prior to transduction. On DIV 11, media was removed and RNA extraction either occurred immediately or the culture plate was stored at −80ºC for RNA extraction at a later date. For the *Reln* targeting experiment, data is shown from neurons that received media supplementation (10μL Neurobasal media) 1hr prior to RNA extraction.

### HEK293T Cell Line

HEK293T cells were obtained from American Type Culture Collection (ATCC catalog #CRL-3216, RRID:CVCL_0063) and were maintained in standard HEK Media: DMEM (DMEM High glucose, pyruvate; Gibco 11995081) + 10% FBS (Qualified US Origin; BioFluid 200-500-Q) + 1U Penicillin-Streptomycin (Gibco 15140122). Cells were passaged in T75 or T225 tissue culture flasks at 70-80% confluence no more than 25 times and were checked for mycoplasma contamination periodically. HEK293T cells were utilized in luciferase assay experiments and for lentiviral production.

### Plasmid Design and Construction

To generate a lentivirus-compatible dCas9-KRAB-MeCP2 construct capable of robust neuronal expression, a Gibson assembly cloning strategy was performed using the XhoI and EcoRI restriction sites in our previously published lenti SYN-FLAG-dCas9-VPR backbone (Addgene plasmid #114196^44^), substituting VPR for KRAB-MeCP2 via PCR amplification from the original dCas9-KRAB-MeCP2 vector (a gift from Alejandro Chavez & George Church (Addgene plasmid #110821^45^)). The full sequence of the updated lentivirus compatible SYN-dCas9-KRAB-MeCP2 is provided in **Supplementary Material**.

A *Fos*-Firefly Luciferase reporter plasmid was generated via amplification of a portion of the *Fos* promoter (−722bp to +97bp of the Rn6 annotated *Fos* transcription start site) from rat genomic DNA. PCR amplification occurred using the forward primer tgctagt*ggatcc*TTGTAGGTAAAGCGGGTTATTGA and reverse primer tgctagt*aagctt*GGGTAGACACTGGTGGGA, followed by a BamHI or HindIII digestion and ligation reaction to insert this sequence upstream of a firefly luciferase construct (a gift from Michael Rehli). The full sequence of the *Fos*-Firefly Luciferase reporter construct is found in **Supplementary Material**.

All sgRNAs utilized here were cloned into our previously published lenti U6-sgRNA/EF1α-mCherry vector using BbsI restriction digest and sense/antisense oligos containing the target sequence, as previously described (Addgene plasmid #114199^44,46^). All sgRNAs were designed using the online tool ChopChop (http://chopchop.cgu.uib.no)^51^, and examined for genome-wide sequence specificity using the National Center for Biotechnology Information’s (NCBI) Basic Local Alignment Search Tool (BLAST). shRNA construction followed a similar approach, utilizing the Broad TRC shRNA design tool (http://portals.broadinstitute.org/gpp/public/) and inserted into a lenti U6-shRNA/EF1α-mCherry vector^52^. shRNAs were assessed for specificity using BLAST. A list of all sgRNA and shRNA target sequences is provided in **Supplementary Table 1**.

### Luciferase Assay

80,000 HEK293T cells were plated in 500μl HEK Media and 24 hrs later 500ng total plasmid DNA was transfected with 1.5μl FuGene HD (Promega) as follows: 50ng of luciferase plasmid, 450ng in 1:2 molar ratio of total sgRNA:dCas9-KRAB-MeCP2. 24hrs following transfection, a luciferase glow assay was performed according to manufacturer’s instructions (Thermo Scientific Pierce Firefly Glow Assay; Thermo Scientific 16177). Briefly, cells were lysed in 200μl 1× Cell Lysis Buffer while shaking at low speed and protected from light for 45 min. 20μl of lysate was then added to an opaque 96-well microplate (Corning 353296) and combined with 50μl 1× D-Luciferin Working Solution supplemented with 1× Firefly Signal Enhancer (Thermo Scientific Pierce Firefly Signal Enhancer; Thermo Scientific 16180). Following a 10 min dark incubation period to allow for signal stabilization, luminescence was recorded using a Synergy 2 Multi-Detection Microplate Reader (BioTek) with a read height of 1mm, a 1 sec integration time, and a 100 msec delay.

### Lentivirus Production

Packaging and concentration of the dCas9-KRAB-MeCP2, sgRNA, and shRNA constructs into lentiviruses occurred as described previously^44^. Under sterile BSL-2 conditions, HEK293T cells were transfected with either the dCas9-KRAB-MeCP2, sgRNA, or shRNA constructs in combination with the psPAX2 packaging and pCMV-VSV-G envelope plasmids (Addgene plasmid #12260 and #8454) using FuGene HD (Promega) in fresh HEK media. 48 hrs following transfection, supernatant was removed, large debris were removed by a 10 min spin at 2300 rcf, followed by filtration through a 0.45μm filter, and centrifugation for 1 hr 45 mins at 106,883 rcf at 4ºC. The viral pellet was allowed to resuspend in sterile PBS at 4ºC overnight and stored at −80ºC. Genomic titer was determined using the Lenti-Z qRT-PCR Titration Kit according to manufacturer’s instructions (Takara #631235). Smaller scale virus prep for sgRNAs and shRNAs was performed through a similar transfection in a 12-well culture plate. After 48 hrs lentiviruses were concentrated with Lenti-X concentrator (Takara), resuspended in sterile PBS over 24-48 hrs and used immediately. dCas9-KRAB-MeCP2 lentivirus was used at a genomic titer of >2000 genomic copies/cell unless otherwise indicated in combination with sgRNAs either concentrated to >1000 genomic copies/cell or produced in small scale prep as indicated above and divided across 3 culture plate wells. shRNA-expressing lentiviruses were produced using the small-scale prep described above and divided across 2 culture well per experiment.

### Immunocytochemistry

After removal of neuronal culture media, cells were washed with PBS and incubated at room temperature for 20 min in freshly prepared 4% paraformaldehyde in PBS. After fixation, cells were washed twice with PBS and neuronal membranes were permeabilized with PBS containing 0.25% Triton X-100 for 15 min at room temperature. Cells were then washed three times in PBS, blocked for 1 hr (10% Thermo Blocker bovine serum albumin (BSA) #37525, 0.05% Tween-20, and 300 mM glycine in PBS) and co-incubated with DYDDDDK Tag (FLAG epitope) Monoclonal Antibody (FG4R) (1:5000 in PBS with 10% Thermo Blocker BSA; Thermo Fisher catalog #MA1-91878, RRID: AB_1957945) overnight at 4°C. Cells were then washed three times with PBS before a 45 min incubation in IRDye 680RD Goat anti-Mouse IgG Secondary Antibody (1:250 in PBS with 10% Thermo Blocker BSA; Li-Cor catalog #925-68070, RRID: AB_2651128). Cells were then washed a final three times with PBS for 5 min. Slide covers slips with Prolong Gold anti-fade medium (Invitrogen) containing 4,6-diamidino-2-phenylindole (DAPI) stain were placed atop the culture wells. A Nikon TiS inverted fluorescent microscope was used to capture 10× magnification (1,888mm^2^ field of view) images from a 24-well culture plate.

### RNA extraction and RT-qPCR

Total RNA was extracted (RNAeasy kit, Qiagen) and reverse-transcribed (iScript cDNA Synthesis Kit, Bio-Rad) following the manufacturers’ instructions. cDNA was subject to RT-qPCR for genes of interest in duplicate using a CFX96 real-time PCR system (Bio-Rad) at 95 °C for 3 min, followed by 40 cycles of 95 °C for 10 s and 58 °C for 30 s, followed by real-time melt analysis to verify product specificity, as described previously^44,49,50^. *Gapdh* was used for normalization via the ΔΔCt method^53^. A list of PCR primer sequences is provided in **Supplementary Table 1**.

### Statistical Analysis

Gene expression differences from RT-qPCR experiments were compared with either an unpaired *t*-test or one- or two-way ANOVA with Tukey’s *post-hoc* test where appropriate. Statistical significance was designated at α = 0.05 for all analyses. Statistical and graphical analyses were performed with Prism software (GraphPad). Statistical assumptions (e.g., normality and homogeneity for parametric tests) were formally tested and examined via boxplots.

## Construct and Data Availability

The SYN-dCas9-KRAB-MeCP2 construct generated in this study will be made available on Addgene for use by the broader research community. All relevant data that support the findings of this study are available by request from the corresponding author.

## Author Contributions

C.G.D. conceived of the SYN-dCas9-KRAB-MeCP2 construct, which was created with a cloning strategy designed by J.S.R. with assistance from N.T.S. and N.V.N.C. F.A.S. conceived, designed, and constructed the *Fos*-luciferase construct, and the general shRNA and sgRNA backbones. C.G.D. completed and analyzed the luciferase assay and ICC experiments. S.V.B. designed and constructed the *Bdnf* sgRNA and shRNA constructs and completed and analyzed all *Bdnf* and *Fosb* experiments. N.D. completed and analyzed the *Reln* targeting experiment with assistance from R.A.P. C.G.D. completed and analyzed the *Kmt2b* and *Npy* targeting experiment. N.V.N.C. designed and constructed the *Kmt2b* sgRNA constructs. *Npy* sgRNAs were designed by J.J.D. and constructed by A.J.B. All projects were supervised by J.J.D. C.G.D. and J.J.D. wrote the main text of the manuscript. All authors have approved the final version of the manuscript.

## Acknowledgements

This work was supported by NIH grants DP1-DA039650, R00-DA034681, and R01-MH114990 (J.J.D.), F32-MH112304 (S.V.B.), F32-DA041778 (F.A.S.), T32-GM008361, and T32-NS061788 (C.G.D.). Additional assistance to J.J.D. was provided by the UAB Pittman Scholars Program. We thank all current and former Day Lab members for assistance and support.

