## Supplementary Material for "An improved CRISPR/dCas9 interference tool for neuronal gene suppression"

**lenti SYN-FLAG-dCas9-KRAB-MeCP2 vector sequence:**

SYN Promoter Start-Codon 3xFLAG NLS dCas9 KRAB MeCP2 Stop-Codon WPRE

agtgcaagtgggttttaggaccaggatgaggcggggtgggggtgcctacctgacgaccgaccccgacccactggacaagcacccaacccccattccccaaattgcgcatcccctatcagagagggggaggggaaacaggatgcggcgaggcgcgtgcgcactgccagcttcagcaccgcggacagtgccttcgcccccgcctggcggcgcgcgccaccgccgcctcagcactgaaggcgcgctgacgtcactcgccggtcccccgcaaactccccttcccggccaccttggtcgcgtccgcgccgccgccggcccagccggaccgcaccacgcgaggcgcgagataggggggcacgggcgcgaccatctgcgctgcggcgccggcgactcagcgctgcctcagtctgcggtgggcagcggaggagtcgtgtcgtgcctgagagcgcagACCGGTGCCACCATGGACTATAAGGACCACGACGGAGACTACAAGGATCATGATATTGATTACAAAGACGATGACGATAAGATGGCCCCAAAGAAGAAGCGGAAGGTCGGTATCCACGGAGTCCCAGCAGCCGACAAGAAGTACAGCATCGGCCTGGCCATCGGCACCAACTCTGTGGGCTGGGCCGTGATCACCGACGAGTACAAGGTGCCCAGCAAGAAATTCAAGGTGCTGGGCAACACCGACCGGCACAGCATCAAGAAGAACCTGATCGGAGCCCTGCTGTTCGACAGCGGCGAAACAGCCGAGGCCACCCGGCTGAAGAGAACCGCCAGAAGAAGATACACCAGACGGAAGAACCGGATCTGCTATCTGCAAGAGATCTTCAGCAACGAGATGGCCAAGGTGGACGACAGCTTCTTCCACAGACTGGAAGAGTCCTTCCTGGTGGAAGAGGATAAGAAGCACGAGCGGCACCCCATCTTCGGCAACATCGTGGACGAGGTGGCCTACCACGAGAAGTACCCCACCATCTACCACCTGAGAAAGAAACTGGTGGACAGCACCGACAAGGCCGACCTGCGGCTGATCTATCTGGCCCTGGCCCACATGATCAAGTTCCGGGGCCACTTCCTGATCGAGGGCGACCTGAACCCCGACAACAGCGACGTGGACAAGCTGTTCATCCAGCTGGTGCAGACCTACAACCAGCTGTTCGAGGAAAACCCCATCAACGCCAGCGGCGTGGACGCCAAGGCCATCCTGTCTGCCAGACTGAGCAAGAGCAGACGGCTGGAAAATCTGATCGCCCAGCTGCCCGGCGAGAAGAAGAATGGCCTGTTCGGCAACCTGATTGCCCTGAGCCTGGGCCTGACCCCCAACTTCAAGAGCAACTTCGACCTGGCCGAGGATGCCAAACTGCAGCTGAGCAAGGACACCTACGACGACGACCTGGACAACCTGCTGGCCCAGATCGGCGACCAGTACGCCGACCTGTTTCTGGCCGCCAAGAACCTGTCCGACGCCATCCTGCTGAGCGACATCCTGAGAGTGAACACCGAGATCACCAAGGCCCCCCTGAGCGCCTCTATGATCAAGAGATACGACGAGCACCACCAGGACCTGACCCTGCTGAAAGCTCTCGTGCGGCAGCAGCTGCCTGAGAAGTACAAAGAGATTTTCTTCGACCAGAGCAAGAACGGCTACGCCGGCTACATTGACGGCGGAGCCAGCCAGGAAGAGTTCTACAAGTTCATCAAGCCCATCCTGGAAAAGATGGACGGCACCGAGGAACTGCTCGTGAAGCTGAACAGAGAGGACCTGCTGCGGAAGCAGCGGACCTTCGACAACGGCAGCATCCCCCACCAGATCCACCTGGGAGAGCTGCACGCCATTCTGCGGCGGCAGGAAGATTTTTACCCATTCCTGAAGGACAACCGGGAAAAGATCGAGAAGATCCTGACCTTCCGCATCCCCTACTACGTGGGCCCTCTGGCCAGGGGAAACAGCAGATTCGCCTGGATGACCAGAAAGAGCGAGGAAACCATCACCCCCTGGAACTTCGAGGAAGTGGTGGACAAGGGCGCTTCCGCCCAGAGCTTCATCGAGCGGATGACCAACTTCGATAAGAACCTGCCCAACGAGAAGGTGCTGCCCAAGCACAGCCTGCTGTACGAGTACTTCACCGTGTATAACGAGCTGACCAAAGTGAAATACGTGACCGAGGGAATGAGAAAGCCCGCCTTCCTGAGCGGCGAGCAGAAAAAGGCCATCGTGGACCTGCTGTTCAAGACCAACCGGAAAGTGACCGTGAAGCAGCTGAAAGAGGACTACTTCAAGAAAATCGAGTGCTTCGACTCCGTGGAAATCTCCGGCGTGGAAGATCGGTTCAACGCCTCCCTGGGCACATACCACGATCTGCTGAAAATTATCAAGGACAAGGACTTCCTGGACAATGAGGAAAACGAGGACATTCTGGAAGATATCGTGCTGACCCTGACACTGTTTGAGGACAGAGAGATGATCGAGGAACGGCTGAAAACCTATGCCCACCTGTTCGACGACAAAGTGATGAAGCAGCTGAAGCGGCGGAGATACACCGGCTGGGGCAGGCTGAGCCGGAAGCTGATCAACGGCATCCGGGACAAGCAGTCCGGCAAGACAATCCTGGATTTCCTGAAGTCCGACGGCTTCGCCAACAGAAACTTCATGCAGCTGATCCACGACGACAGCCTGACCTTTAAAGAGGACATCCAGAAAGCCCAGGTGTCCGGCCAGGGCGATAGCCTGCACGAGCACATTGCCAATCTGGCCGGCAGCCCCGCCATTAAGAAGGGCATCCTGCAGACAGTGAAGGTGGTGGACGAGCTCGTGAAAGTGATGGGCCGGCACAAGCCCGAGAACATCGTGATCGAAATGGCCAGAGAGAACCAGACCACCCAGAAGGGACAGAAGAACAGCCGCGAGAGAATGAAGCGGATCGAAGAGGGCATCAAAGAGCTGGGCAGCCAGATCCTGAAAGAACACCCCGTGGAAAACACCCAGCTGCAGAACGAGAAGCTGTACCTGTACTACCTGCAGAATGGGCGGGATATGTACGTGGACCAGGAACTGGACATCAACCGGCTGTCCGACTACGATGTGGACGCTATCGTGCCTCAGAGCTTTCTGAAGGACGACTCCATCGACAACAAGGTGCTGACCAGAAGCGACAAGAACCGGGGCAAGAGCGACAACGTGCCCTCCGAAGAGGTCGTGAAGAAGATGAAGAACTACTGGCGGCAGCTGCTGAACGCCAAGCTGATTACCCAGAGAAAGTTCGACAATCTGACCAAGGCCGAGAGAGGCGGCCTGAGCGAACTGGATAAGGCCGGCTTCATCAAGAGACAGCTGGTGGAAACCCGGCAGATCACAAAGCACGTGGCACAGATCCTGGACTCCCGGATGAACACTAAGTACGACGAGAATGACAAGCTGATCCGGGAAGTGAAAGTGATCACCCTGAAGTCCAAGCTGGTGTCCGATTTCCGGAAGGATTTCCAGTTTTACAAAGTGCGCGAGATCAACAACTACCACCACGCCCACGACGCCTACCTGAACGCCGTCGTGGGAACCGCCCTGATCAAAAAGTACCCTAAGCTGGAAAGCGAGTTCGTGTACGGCGACTACAAGGTGTACGACGTGCGGAAGATGATCGCCAAGAGCGAGCAGGAAATCGGCAAGGCTACCGCCAAGTACTTCTTCTACAGCAACATCATGAACTTTTTCAAGACCGAGATTACCCTGGCCAACGGCGAGATCCGGAAGCGGCCTCTGATCGAGACAAACGGCGAAACCGGGGAGATCGTGTGGGATAAGGGCCGGGATTTTGCCACCGTGCGGAAAGTGCTGAGCATGCCCCAAGTGAATATCGTGAAAAAGACCGAGGTGCAGACAGGCGGCTTCAGCAAAGAGTCTATCCTGCCCAAGAGGAACAGCGATAAGCTGATCGCCAGAAAGAAGGACTGGGACCCTAAGAAGTACGGCGGCTTCGACAGCCCCACCGTGGCCTATTCTGTGCTGGTGGTGGCCAAAGTGGAAAAGGGCAAGTCCAAGAAACTGAAGAGTGTGAAAGAGCTGCTGGGGATCACCATCATGGAAAGAAGCAGCTTCGAGAAGAATCCCATCGACTTTCTGGAAGCCAAGGGCTACAAAGAAGTGAAAAAGGACCTGATCATCAAGCTGCCTAAGTACTCCCTGTTCGAGCTGGAAAACGGCCGGAAGAGAATGCTGGCCTCTGCCGGCGAACTGCAGAAGGGAAACGAACTGGCCCTGCCCTCCAAATATGTGAACTTCCTGTACCTGGCCAGCCACTATGAGAAGCTGAAGGGCTCCCCCGAGGATAATGAGCAGAAACAGCTGTTTGTGGAACAGCACAAGCACTACCTGGACGAGATCATCGAGCAGATCAGCGAGTTCTCCAAGAGAGTGATCCTGGCCGACGCTAATCTGGACAAAGTGCTGTCCGCCTACAACAAGCACCGGGATAAGCCCATCAGAGAGCAGGCCGAGAATATCATCCACCTGTTTACCCTGACCAATCTGGGAGCCCCTGCCGCCTTCAAGTACTTTGACACCACCATCGACCGGAAGAGGTACACCAGCACCAAAGAGGTGCTGGACGCCACCCTGATCCACCAGAGCATCACCGGCCTGTACGAGACACGGATCGACCTGTCTCAGCTGGGAGGCGACAAAAGGCCGGCGGCCACGAAAAAGGCCGGACAGGCCAAAAAGAAAAAGagtggtggaggaagtggcgggtcagggtcgatggacgcgaaatcacttacggcatggtcgagaacactggttacgttcaaggacgtgtttgtggactttacacgtgaggagtggaaattgctggatactgcgcaacaaattgtgtatcgaaatgtcatgcttgagaattacaagaacctcgtcagtctcggataccagttgacgaaaccggatgtgatccttaggctcgaaaagggggaagaaccttggctggtatcgggaggtggttcgggtggctctggatcaagcccaaagaagaaacggaaggtggaagcctcagtgcaggtgaaaagggtgctggaaaaatcccccggcaaactcctcgtgaagatgcccttccaggcttcccctggcggaaaaggtgaagggggtggcgcaaccacatctgcccaggtcatggtcatcaagcgacctggaaggaaaagaaaggccgaggctgaccctcaggccattccaaagaaacggggacgcaagccagggtccgtggtcgcagctgcagcagctgaggctaagaaaaaggcagtgaaggaaagctccatccgcagtgtgcaggagactgtcctgcccatcaagaagaggaagactagggagaccgtgtccatcgaggtcaaagaagtggtcaagcccctgctcgtgtccaccctgggcgaaaaatctggaaaggggctcaaaacatgcaagtcacctggacggaaaagcaaggagtctagtccaaaggggcgctcaagctccgcttctagtccccctaaaaaggaacaccatcaccatcaccatcacgccgagtctcctaaggctcctatgccactgctcccaccacctccaccacctgagccacagtcaagcgaagaccccatcagcccacccgagcctcaggatctgtcctctagtatttgcaaagaggaaaagatgcccagagcaggcagcctggagagtgatggctgtccaaaagaacccgccaagacccagcctatggtggcagccgctgcaactaccaccacaaccacaactaccacagtggccgaaaaatacaagcatcgcggcgagggcgaacgaaaggacattgtgtcaagctccatgcccagacctaaccgggaggaaccagtcgatagtaggacacccgtgactgagagagtctcatagGAATTCAATTCTAACTAGACGCGTTAAGTCGACAATCAACCTCTGGATTACAAAATTTGTGAAAGATTGACTGGTATTCTTAACTATGTTGCTCCTTTTACGCTATGTGGATACGCTGCTTTAATGCCTTTGTATCATGCTATTGCTTCCCGTATGGCTTTCATTTTCTCCTCCTTGTATAAATCCTGGTTGCTGTCTCTTTATGAGGAGTTGTGGCCCGTTGTCAGGCAACGTGGCGTGGTGTGCACTGTGTTTGCTGACGCAACCCCCACTGGTTGGGGCATTGCCACCACCTGTCAGCTCCTTTCCGGGACTTTCGCTTTCCCCCTCCCTATTGCCACGGCGGAACTCATCGCCGCCTGCCTTGCCCGCTGCTGGACAGGGGCTCGGCTGTTGGGCACTGACAATTCCGTGGTGTTGTCGGGGAAATCATCGTCCTTTCCTTGGCTGCTCGCCTGTGTTGCCACCTGGATTCTGCGCGGGACGTCCTTCTGCTACGTCCCTTCGGCCCTCAATCCAGCGGACCTTCCTTCCCGCGGCCTGCTGCCGGCTCTGCGGCCTCTTCCGCGTCTTCGCCTTCGCCCTCAGACGAGTCGGATCTCCCTTTGGGCCGCCTCCCCGCGTCGACTTTAAGACCAATGACTTACAAGGCAGCTGTAGATCTTAGCCACTTTTTAAAAGAAAAGGGGGGACTGGAAGGGCTAATTCACTCCCAACGAAGACAAGATCTGCTTTTTGCTTGTACTGGGTCTCTCTGGTTAGACCAGATCTGAGCCTGGGAGCTCTCTGGCTAACTAGGGAACCCACTGCTTAAGCCTCAATAAAGCTTGCCTTGAGTGCTTCAAGTAGTGTGTGCCCGTCTGTTGTGTGACTCTGGTAACTAGAGATCCCTCAGACCCTTTTAGTCAGTGTGGAAAATCTCTAGCAGGGCCCGTTTAAACCCGCTGATCAGCCTCGACTGTGCCTTCTAGTTGCCAGCCATCTGTTGTTTGCCCCTCCCCCGTGCCTTCCTTGACCCTGGAAGGTGCCACTCCCACTGTCCTTTCCTAATAAAATGAGGAAATTGCATCGCATTGTCTGAGTAGGTGTCATTCTATTCTGGGGGGTGGGGTGGGGCAGGACAGCAAGGGGGAGGATTGGGAAGACAATAGCAGGCATGCTGGGGATGCGGTGGGCTCTATGGCTTCTGAGGCGGAAAGAACCAGCTGGGGCTCTAGGGGGTATCCCCACGCGCCCTGTAGCGGCGCATTAAGCGCGGCGGGTGTGGTGGTTACGCGCAGCGTGACCGCTACACTTGCCAGCGCCCTAGCGCCCGCTCCTTTCGCTTTCTTCCCTTCCTTTCTCGCCACGTTCGCCGGCTTTCCCCGTCAAGCTCTAAATCGGGGGCTCCCTTTAGGGTTCCGATTTAGTGCTTTACGGCACCTCGACCCCAAAAAACTTGATTAGGGTGATGGTTCACGTAGTGGGCCATCGCCCTGATAGACGGTTTTTCGCCCTTTGACGTTGGAGTCCACGTTCTTTAATAGTGGACTCTTGTTCCAAACTGGAACAACACTCAACCCTATCTCGGTCTATTCTTTTGATTTATAAGGGATTTTGCCGATTTCGGCCTATTGGTTAAAAAATGAGCTGATTTAACAAAAATTTAACGCGAATTAATTCTGTGGAATGTGTGTCAGTTAGGGTGTGGAAAGTCCCCAGGCTCCCCAGCAGGCAGAAGTATGCAAAGCATGCATCTCAATTAGTCAGCAACCAGGTGTGGAAAGTCCCCAGGCTCCCCAGCAGGCAGAAGTATGCAAAGCATGCATCTCAATTAGTCAGCAACCATAGTCCCGCCCCTAACTCCGCCCATCCCGCCCCTAACTCCGCCCAGTTCCGCCCATTCTCCGCCCCATGGCTGACTAATTTTTTTTATTTATGCAGAGGCCGAGGCCGCCTCTGCCTCTGAGCTATTCCAGAAGTAGTGAGGAGGCTTTTTTGGAGGCCTAGGCTTTTGCAAAAAGCTCCCGGGAGCTTGTATATCCATTTTCGGATCTGATCAGCACGTGTTGACAATTAATCATCGGCATAGTATATCGGCATAGTATAATACGACAAGGTGAGGAACTAAACCATGGCCAAGTTGACCAGTGCCGTTCCGGTGCTCACCGCGCGCGACGTCGCCGGAGCGGTCGAGTTCTGGACCGACCGGCTCGGGTTCTCCCGGGACTTCGTGGAGGACGACTTCGCCGGTGTGGTCCGGGACGACGTGACCCTGTTCATCAGCGCGGTCCAGGACCAGGTGGTGCCGGACAACACCCTGGCCTGGGTGTGGGTGCGCGGCCTGGACGAGCTGTACGCCGAGTGGTCGGAGGTCGTGTCCACGAACTTCCGGGACGCCTCCGGGCCGGCCATGACCGAGATCGGCGAGCAGCCGTGGGGGCGGGAGTTCGCCCTGCGCGACCCGGCCGGCAACTGCGTGCACTTCGTGGCCGAGGAGCAGGACTGACACGTGCTACGAGATTTCGATTCCACCGCCGCCTTCTATGAAAGGTTGGGCTTCGGAATCGTTTTCCGGGACGCCGGCTGGATGATCCTCCAGCGCGGGGATCTCATGCTGGAGTTCTTCGCCCACCCCAACTTGTTTATTGCAGCTTATAATGGTTACAAATAAAGCAATAGCATCACAAATTTCACAAATAAAGCATTTTTTTCACTGCATTCTAGTTGTGGTTTGTCCAAACTCATCAATGTATCTTATCATGTCTGTATACCGTCGACCTCTAGCTAGAGCTTGGCGTAATCATGGTCATAGCTGTTTCCTGTGTGAAATTGTTATCCGCTCACAATTCCACACAACATACGAGCCGGAAGCATAAAGTGTAAAGCCTGGGGTGCCTAATGAGTGAGCTAACTCACATTAATTGCGTTGCGCTCACTGCCCGCTTTCCAGTCGGGAAACCTGTCGTGCCAGCTGCATTAATGAATCGGCCAACGCGCGGGGAGAGGCGGTTTGCGTATTGGGCGCTCTTCCGCTTCCTCGCTCACTGACTCGCTGCGCTCGGTCGTTCGGCTGCGGCGAGCGGTATCAGCTCACTCAAAGGCGGTAATACGGTTATCCACAGAATCAGGGGATAACGCAGGAAAGAACATGTGAGCAAAAGGCCAGCAAAAGGCCAGGAACCGTAAAAAGGCCGCGTTGCTGGCGTTTTTCCATAGGCTCCGCCCCCCTGACGAGCATCACAAAAATCGACGCTCAAGTCAGAGGTGGCGAAACCCGACAGGACTATAAAGATACCAGGCGTTTCCCCCTGGAAGCTCCCTCGTGCGCTCTCCTGTTCCGACCCTGCCGCTTACCGGATACCTGTCCGCCTTTCTCCCTTCGGGAAGCGTGGCGCTTTCTCATAGCTCACGCTGTAGGTATCTCAGTTCGGTGTAGGTCGTTCGCTCCAAGCTGGGCTGTGTGCACGAACCCCCCGTTCAGCCCGACCGCTGCGCCTTATCCGGTAACTATCGTCTTGAGTCCAACCCGGTAAGACACGACTTATCGCCACTGGCAGCAGCCACTGGTAACAGGATTAGCAGAGCGAGGTATGTAGGCGGTGCTACAGAGTTCTTGAAGTGGTGGCCTAACTACGGCTACACTAGAAGAACAGTATTTGGTATCTGCGCTCTGCTGAAGCCAGTTACCTTCGGAAAAAGAGTTGGTAGCTCTTGATCCGGCAAACAAACCACCGCTGGTAGCGGTGGTTTTTTTGTTTGCAAGCAGCAGATTACGCGCAGAAAAAAAGGATCTCAAGAAGATCCTTTGATCTTTTCTACGGGGTCTGACGCTCAGTGGAACGAAAACTCACGTTAAGGGATTTTGGTCATGAGATTATCAAAAAGGATCTTCACCTAGATCCTTTTAAATTAAAAATGAAGTTTTAAATCAATCTAAAGTATATATGAGTAAACTTGGTCTGACAGTTACCAATGCTTAATCAGTGAGGCACCTATCTCAGCGATCTGTCTATTTCGTTCATCCATAGTTGCCTGACTCCCCGTCGTGTAGATAACTACGATACGGGAGGGCTTACCATCTGGCCCCAGTGCTGCAATGATACCGCGAGACCCACGCTCACCGGCTCCAGATTTATCAGCAATAAACCAGCCAGCCGGAAGGGCCGAGCGCAGAAGTGGTCCTGCAACTTTATCCGCCTCCATCCAGTCTATTAATTGTTGCCGGGAAGCTAGAGTAAGTAGTTCGCCAGTTAATAGTTTGCGCAACGTTGTTGCCATTGCTACAGGCATCGTGGTGTCACGCTCGTCGTTTGGTATGGCTTCATTCAGCTCCGGTTCCCAACGATCAAGGCGAGTTACATGATCCCCCATGTTGTGCAAAAAAGCGGTTAGCTCCTTCGGTCCTCCGATCGTTGTCAGAAGTAAGTTGGCCGCAGTGTTATCACTCATGGTTATGGCAGCACTGCATAATTCTCTTACTGTCATGCCATCCGTAAGATGCTTTTCTGTGACTGGTGAGTACTCAACCAAGTCATTCTGAGAATAGTGTATGCGGCGACCGAGTTGCTCTTGCCCGGCGTCAATACGGGATAATACCGCGCCACATAGCAGAACTTTAAAAGTGCTCATCATTGGAAAACGTTCTTCGGGGCGAAAACTCTCAAGGATCTTACCGCTGTTGAGATCCAGTTCGATGTAACCCACTCGTGCACCCAACTGATCTTCAGCATCTTTTACTTTCACCAGCGTTTCTGGGTGAGCAAAAACAGGAAGGCAAAATGCCGCAAAAAAGGGAATAAGGGCGACACGGAAATGTTGAATACTCATACTCTTCCTTTTTCAATATTATTGAAGCATTTATCAGGGTTATTGTCTCATGAGCGGATACATATTTGAATGTATTTAGAAAAATAAACAAATAGGGGTTCCGCGCACATTTCCCCGAAAAGTGCCACCTGACGTCGACGGATCGGGAGATCTCCCGATCCCCTATGGTGCACTCTCAGTACAATCTGCTCTGATGCCGCATAGTTAAGCCAGTATCTGCTCCCTGCTTGTGTGTTGGAGGTCGCTGAGTAGTGCGCGAGCAAAATTTAAGCTACAACAAGGCAAGGCTTGACCGACAATTGCATGAAGAATCTGCTTAGGGTTAGGCGTTTTGCGCTGCTTCGCGATGTACGGGCCAGATATACGCGTTGACATTGATTATTGACTAGTTATTAATAGTAATCAATTACGGGGTCATTAGTTCATAGCCCATATATGGAGTTCCGCGTTACATAACTTACGGTAAATGGCCCGCCTGGCTGACCGCCCAACGACCCCCGCCCATTGACGTCAATAATGACGTATGTTCCCATAGTAACGCCAATAGGGACTTTCCATTGACGTCAATGGGTGGAGTATTTACGGTAAACTGCCCACTTGGCAGTACATCAAGTGTATCATATGCCAAGTACGCCCCCTATTGACGTCAATGACGGTAAATGGCCCGCCTGGCATTATGCCCAGTACATGACCTTATGGGACTTTCCTACTTGGCAGTACATCTACGTATTAGTCATCGCTATTACCATGGTGATGCGGTTTTGGCAGTACATCAATGGGCGTGGATAGCGGTTTGACTCACGGGGATTTCCAAGTCTCCACCCCATTGACGTCAATGGGAGTTTGTTTTGGCACCAAAATCAACGGGACTTTCCAAAATGTCGTAACAACTCCGCCCCATTGACGCAAATGGGCGGTAGGCGTGTACGGTGGGAGGTCTATATAAGCAGCGCGTTTTGCCTGTACTGGGTCTCTCTGGTTAGACCAGATCTGAGCCTGGGAGCTCTCTGGCTAACTAGGGAACCCACTGCTTAAGCCTCAATAAAGCTTGCCTTGAGTGCTTCAAGTAGTGTGTGCCCGTCTGTTGTGTGACTCTGGTAACTAGAGATCCCTCAGACCCTTTTAGTCAGTGTGGAAAATCTCTAGCAGTGGCGCCCGAACAGGGACTTGAAAGCGAAAGGGAAACCAGAGGAGCTCTCTCGACGCAGGACTCGGCTTGCTGAAGCGCGCACGGCAAGAGGCGAGGGGCGGCGACTGGTGAGTACGCCAAAAATTTTGACTAGCGGAGGCTAGAAGGAGAGAGATGGGTGCGAGAGCGTCAGTATTAAGCGGGGGAGAATTAGATCGCGATGGGAAAAAATTCGGTTAAGGCCAGGGGGAAAGAAAAAATATAAATTAAAACATATAGTATGGGCAAGCAGGGAGCTAGAACGATTCGCAGTTAATCCTGGCCTGTTAGAAACATCAGAAGGCTGTAGACAAATACTGGGACAGCTACAACCATCCCTTCAGACAGGATCAGAAGAACTTAGATCATTATATAATACAGTAGCAACCCTCTATTGTGTGCATCAAAGGATAGAGATAAAAGACACCAAGGAAGCTTTAGACAAGATAGAGGAAGAGCAAAACAAAAGTAAGACCACCGCACAGCAAGCGGCCGCTGATCTTCAGACCTGGAGGAGGAGATATGAGGGACAATTGGAGAAGTGAATTATATAAATATAAAGTAGTAAAAATTGAACCATTAGGAGTAGCACCCACCAAGGCAAAGAGAAGAGTGGTGCAGAGAGAAAAAAGAGCAGTGGGAATAGGAGCTTTGTTCCTTGGGTTCTTGGGAGCAGCAGGAAGCACTATGGGCGCAGCGTCAATGACGCTGACGGTACAGGCCAGACAATTATTGTCTGGTATAGTGCAGCAGCAGAACAATTTGCTGAGGGCTATTGAGGCGCAACAGCATCTGTTGCAACTCACAGTCTGGGGCATCAAGCAGCTCCAGGCAAGAATCCTGGCTGTGGAAAGATACCTAAAGGATCAACAGCTCCTGGGGATTTGGGGTTGCTCTGGAAAACTCATTTGCACCACTGCTGTGCCTTGGAATGCTAGTTGGAGTAATAAATCTCTGGAACAGATTTGGAATCACACGACCTGGATGGAGTGGGACAGAGAAATTAACAATTACACAAGCTTAATACACTCCTTAATTGAAGAATCGCAAAACCAGCAAGAAAAGAATGAACAAGAATTATTGGAATTAGATAAATGGGCAAGTTTGTGGAATTGGTTTAACATAACAAATTGGCTGTGGTATATAAAATTATTCATAATGATAGTAGGAGGCTTGGTAGGTTTAAGAATAGTTTTTGCTGTACTTTCTATAGTGAATAGAGTTAGGCAGGGATATTCACCATTATCGTTTCAGACCCACCTCCCAACCCCGAGGGGACCCGACAGGCCCGAAGGAATAGAAGAAGAAGGTGGAGAGAGAGACAGAGACAGATCCATTCGATTAGTGAACGGATCGGCACTGCGTGCGCCAATTCTGCAGACAAATGGCAGTATTCATCCACAATTTTAAAAGAAAAGGGGGGATTGGGGGGTACAGTGCAGGGGAAAGAATAGTAGACATAATAGCAACAGACATACAAACTAAAGAATTACAAAAACAAATTACAAAAATTCAAAATTTTCGGGTTTATTACAGGGACAGCAGAGATCCAGTTTGGTTAATTAA

***Fos*-FireflyLuciferase Reporter:**

*Fos*-Promoter Annotated-*Fos*-TSS Start-Codon Firefly-Luciferase Stop-Codon SV40-PolyA-Signal

ttaattaaaattatctctaaggcatgtgaactggctgtcttggttttcatctgtacttcatctgctacctctgtgacctgaaacatatttataattccattaagctgtgcatatgatagatttatcatatgtattttccttaaaggatttttgtaagaactaattgaattgatacctgtaaagtctttatcacactacccaataaataataaatctctttgttcagctctctgtttctataaatatgtaccagttttattgtttttagtggtagtgattttattctctttctatatatatacacacacatgtgtgcattcataaatatatacaatttttatgaataaaaaattattagcaatcaatattgaaaaccactgatttttgtttatgtgagcaaacagcagattaaaaggaattcctgcaggactagtggatccTTGTAGGTAAAGCGGGTTATTGAATACTTACTTAGAATCCTTCACTTACTGCTTCCAACCTCAGGCCTAATGTTGCACTGATTTGGGACGGAGAGAGGTCTGATGTGGGCTAGCTTTCCTTTGGGAACAGAGGCTTGGAGCCTTTAGGGCTGCGTGCCTGCTTCTCCTAATACCAGAGACTTTTTTAAAAAGCTCCAGATTGCTGGACAATGGAAAGGAGATGACCCCCAGTCTCATCCCCTGACCCTGGGAACAGAGTACACATTGAATCAGGTGCGAATGTTCGCTCGCCTTCTCTGCCTTTCCCGCCTCCCCTCCCCCGGCCGCGGCCCCCGCTCCCCCCTTGCGCTGCACCCTCAGAGTTGGCTGCAGCCGGCGAGCTGTTCCCGTCAATCCCTCCCTCCTTTACACAGGATGTCCATATTAGGACATCTGCGTCAGCAGGTTTCCACGGCCGGTCCCTGTTGTCCTGGGGGGAACCATCCCCGAAATCCTACATGCGGAGGGTCCAGGAGACCTTCTAAGATCCCAATTGTGAACACTCATAGGTGAAAGTTACAGACTGAGACGGGGGTTGAGAGCCTGGGGCGTAGAGTTGATGACAGGGAGCCCGCAGAGGGCATTCGGGAGCGCTTTCCCCCCTCCAGTTTCTCTGTTCCGCTCATGACGTAGTAAGCCATTCAAGCGCTTCTATAAAGCGGCCAGCTGAGGCGCCTACTACTCCAACCGCGATTGCAGCTAGCAACTGAGAAGACTGGATAGAGCCGGCGGAGCCGCGAACGAGCAGTGACCGCGCTCCCACCCAGCTCTGCTCTGCAGCTCCCACCAGTGTCTACCCaagcttagtccatggaggatgccaagaatattaagaaaggccctgccccattctaccctctggaagatggcactgctggtgagcaactgcacaaggccatgaagaggtatgccctggtccctggcaccattgccttcactgatgctcacattgaggtggacatcacctatgctgaatactttgagatgtctgtgaggctggcagaagccatgaaaagatatggactgaacaccaaccacaggattgtggtgtgctctgagaactctctccagttcttcatgcctgtgttaggagccctgttcattggagtggctgtggcccctgccaatgacatctacaatgagagagagctcctgaacagcatgggcatcagccagccaactgtggtctttgtgagcaagaagggcctgcaaaagatcctgaatgtgcagaagaagctgcccatcatccagaagatcatcatcatggacagcaagactgactaccagggcttccagagcatgtatacctttgtgaccagccacttaccccctggcttcaatgagtatgactttgtgcctgagagctttgacagggacaagaccattgctctgattatgaacagctctggctccactggactgcccaaaggtgtggctctgccccacagaactgcttgtgtgagattcagccatgccagagaccccatctttggcaaccagatcatccctgacactgccatcctgtctgtggttccattccatcatggctttggcatgttcacaacactggggtacctgatctgtggcttcagagtggtgctgatgtataggtttgaggaggagctgtttctgaggagcctacaagactacaagatccagtctgccctgctggtgcccactctgttcagcttctttgccaagagcaccctcattgacaagtatgacctgagcaacctgcatgagattgcctctggaggagcacccctgagcaaggaggtgggtgaggctgtggcaaagaggttccatctcccaggaatcagacagggctatggcctgactgagaccacctctgccatcctcatcacccctgaaggagatgacaagcctggtgctgtgggcaaggtggttcccttttttgaggccaaggtggtggacctggacactggcaagaccctgggagtgaaccagaggggtgagctgtgtgtgaggggtcccatgatcatgtctggctatgtgaacaaccctgaggccaccaatgccctgattgacaaggatggctggctgcactctggtgacattgcctactgggatgaggatgagcactttttcattgtggacaggctgaagagcctcatcaagtacaaaggctaccaagtggcacctgctgagctagagagcatcctgctccagcaccccaacatctttgatgctggtgtggctggcctgcctgatgatgatgctggagagctgcctgctgctgttgtggttctggagcatggaaagaccatgactgagaaggagattgtggactatgtggccagtcaggtgaccactgccaagaagctgaggggaggtgtggtgtttgtggatgaggtgccaaagggtctgactggcaagctggatgccagaaagatcagagagatcctgatcaaggccaagaagggtggcaaacaattctagctggccagacatgataagatacattgatgagtttggacaaaccacaactagaatgcagtgaaaaaaatgctttatttgtgaaatttgtgatgctattgctttatttgtaaccattataagctgcaataaacaagttaacaacaacaattgcattcattttatgtttcaggttcagggggaggtgtgggaggttttttaaagcaagtaaaacctctacaaatgtggtatggaattcagtcaatatgttcaccccaaaaaagctgtttgttaacttgtcaacctcattctaaaatgtatatagaagcccaaaagacaataacaaaaatattcttgtagaacaaaatgggaaagaatgttccactaaatatcaagatttagagcaaagcatgagatgtgtggggatagacagtgaggctgataaaatagagtagagctcagaaacagacccattgatatatgtaagtgacctatgaaaaaaatatggcattttacaatgggaaaatgatgatctttttcttttttagaaaaacagggaaatatatttatatgtaaaaaataaaagggaacccatatgtcataccatacacacaaaaaaattccagtgaattataagtctaaatggagaaggcaaaactttaaatcttttagaaaataatatagaagcatgccatcaagacttcagtgtagagaaaaatttcttatgactcaaagtcctaaccacaaagaaaagattgttaattagattgcatgaatattaagacttatttttaaaattaaaaaaccattaagaaaagtcaggccatagaatgacagaaaatatttgcaacaccccagtaaagagaattgtaatatgcagattataaaaagaagtcttacaaatcagtaaaaaataaaactagacaaaaatttgaacagatgaaagagaaactctaaataatcattacacatgagaaactcaatctcagaaatcagagaactatcattgcatatacactaaattagagaaatattaaaaggctaagtaacatctgtggcttaattaaaacagtagttgacaattaaacattggcatagtatatctgcatagtataatacaactcactataggagggccatcatggccaagttgaccagtgctgtcccagtgctcacagccagggatgtggctggagctgttgagttctggactgacaggttggggttctccagagattttgtggaggatgactttgcaggtgtggtcagagatgatgtcaccctgttcatctcagcagtccaggaccaggtggtgcctgacaacaccctggcttgggtgtgggtgagaggactggatgagctgtatgctgagtggagtgaggtggtctccaccaacttcagggatgccagtggccctgccatgacagagattggagagcagccctgggggagagagtttgccctgagagacccagcaggcaactgtgtgcactttgtggcagaggagcaggactgaggataacctaggaaaccttaaaacctttaaaagccttatatattcttttttttcttataaaacttaaaaccttagaggctatttaagttgctgatttatattaattttattgttcaaacatgagagcttagtacatgaaacatgagagcttagtacattagccatgagagcttagtacattagccatgagggtttagttcattaaacatgagagcttagtacattaaacatgagagcttagtacatactatcaacaggttgaactgctgatc
